# Climate change favours African malaria vector mosquitoes

**DOI:** 10.1101/2025.03.17.643604

**Authors:** Tiem van der Deure, David Nogués-Bravo, Lembris Laanyuni Njotto, Anna-Sofie Stensgaard

## Abstract

Malaria, a parasitic disease transmitted by mosquitoes of the genus *Anopheles*, causes half a million deaths annually, mostly among children in Africa. Climate change is expected to significantly alter malaria transmission, but previous forecasts have placed little emphasis on the varying impacts climate change could have on different mosquito vector species. Using extensive mosquito observation datasets and species distribution modelling, we investigate the climatic preferences of six dominant African malaria vector species and how the environmental suitability for these species across sub-Saharan Africa might change due to climate and land use change. We highlight three species for which environmental suitability is consistently associated with higher malaria prevalence and that might be favoured by climate change. Our projections indicate a substantial increase in areas highly suitable for these vectors, underscoring the urgent need to adapt malaria control strategies to shifting vector distributions driven by climate change.

## Main

Malaria is a mosquito-borne parasitic disease with around 250 million cases annually. It causes approximately half a million deaths, more than 90% of which occur in sub-Saharan Africa^1,2^. Malaria transmission is heavily influenced by environmental factors. Warm temperatures accelerate the development of *Plasmodium*, the protozoan parasite that causes malaria, within the mosquito, making transmission more efficient, while rainfall creates breeding sites for mosquitoes, and humidity and temperature affect mosquito survival^3^. As the planet warms, understanding possible shifts in the distribution and burden of malaria is crucial for the continued effectiveness of control and eradication efforts^4,5^. However, despite years of research, the historical and possible future impacts of climate change on malaria have been difficult to resolve^6–8^.

To date, most research on the effects of climate change on malaria has focussed on the temperature dependence of the transmission efficiency of malaria parasites, typically through mechanistic models parameterised on experimental life history data^9–12^. A number of studies suggest climate change could put new human populations at risk in cooler areas^13–16^, in particular in East Africa^17^. In hotter areas such as West Africa and the Sahel, previous modelling has predicted that climate change would not have much impact or even lead to a decline in malaria transmission^7,18–20^.

While these studies have significantly improved our understanding of how temperature affects the malaria parasite’s (*Plasmodium falciparum*) life cycle, they only to a limited extent account for the rather complex ecology and biology of the involved mosquito vectors. Most models focus on a single vector species (usually the archetypical vector *Anopheles gambiae*)^21^, despite the fact that malaria transmission in sub-Saharan Africa involves around 10 dominant and over a dozen secondary vector species^22,23^, each with their unique behaviours, environmental tolerances, and habitat preferences^23^. The diversity of malaria vectors is likely to play a key role in the persistence of malaria transmission, as well as for future transmission patterns. Secondary vectors can elude control measures and sustain residual transmission in response to malaria control efforts^24^. Environmental (e.g. drought^25^) or human (e.g. forest clearance^26^) disruption has for instance been shown to trigger a shift from one malaria vector to another. Potentially, climate change could result in shifts in the relative importance of malaria vectors for malaria transmission, rather than the cessation of transmission.

A growing field of literature is available on the distribution of malaria vectors, including extensive, freely available databases of mosquito observations, as well as distribution maps for nine malaria vector species^22,23,27,28^. How each of these vectors will respond to climatic changes will have major implications for malaria transmission, but surprisingly few studies have focussed on how climate change impacts each of the various mosquito vector species^29^, with a few exceptions^30^.

To address this gap, this study investigates how climate change will shift the distribution of the six most important African malaria vectors and where future hotspots of vector-human overlap may emerge. By combining ecological niche theory^31,32^, species distribution models (SDMs), and machine learning methods^33^ with climate and land-use scenarios, we leverage extensive databases on mosquito occurrences to project future distribution patterns. Understanding these shifts is crucial, as climate change presents a significant challenge to malaria control and eradication efforts^34,35^. By identifying new regions where important malaria vectors might emerge, this study provides critical insights to inform targeted vector control strategies, optimize resource allocation, and inform public health policies^36^.

### Different climatic preferences of malaria vectors

To predict future impacts of climate change on malaria vectors, it is important to first understand differences in their realised climatic niches^37^. Therefore, we first analysed the temperature and precipitation at the occurrence locations of six dominant malaria vector species in Africa. This analysis shows that the investigated mosquito species have markedly different climatic niches (Fig. 1). For instance, *Anopheles arabiensis* and *Anopheles funestus* (Fig. 1b and 1c) are climate generalists with wide climatic niches, while *Anopheles coluzzii* and *Anopheles moucheti* (Fig. 1d and 1e) are constrained to a narrower range of climatic conditions.

**Figure 1.**
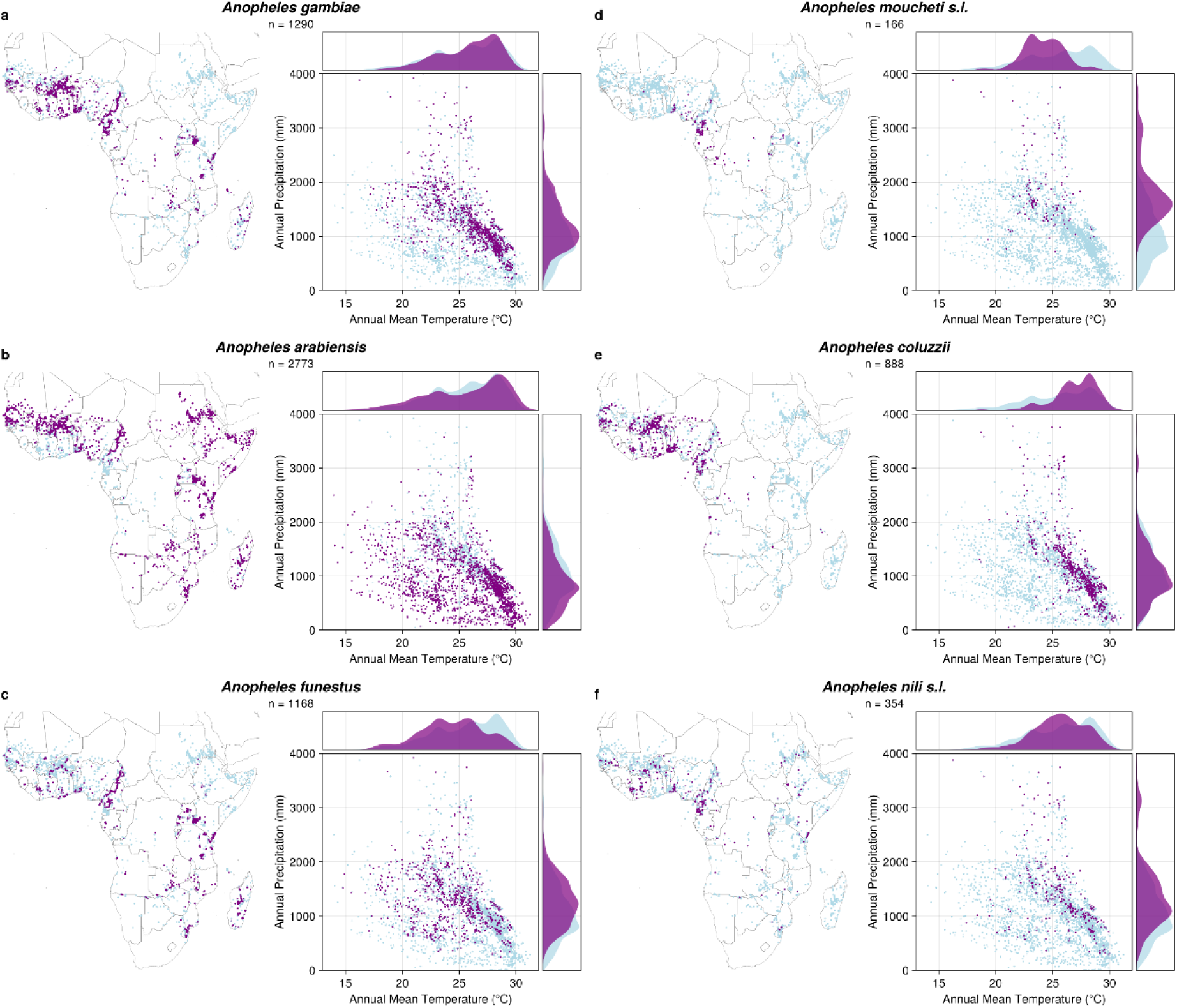
Occurrences records in geographic and climatic space for six important African malaria vectors. For each species, the geographic locations (maps on the left) and climatic conditions (scatter and density plots on the right) are shown in purple, with records of all *Anopheles* species in light blue. The number of records after data cleaning and thinning is also shown.

The realised niches of each species are defined by different temperature and precipitation thresholds. *An. gambiae*, *An. arabiensis*, and *An. coluzzii* are often found in areas with annual mean temperatures of 25°C or above, but while *An. arabiensis* is recorded in extremely dry areas, *An. gambiae* is rarely found in areas with less than 500 mm precipitation per year (Fig. 1). Geographically, *An. arabiensis* is also very widely spread, extending north into the Sahara Desert and south into South Africa. Most records for *An. nili s.l.* have high precipitation, but more moderate temperatures than for *An. gambiae*. *An. moucheti s.l.* is the most specialised of all investigated species and is exclusively recorded in tropical areas in Central Africa, with intermediate temperatures (around 25°C) and more than 1000 mm annual precipitation. Finally, *An. funestus* is relatively often found in East Africa, including in more temperate areas with intermediate temperatures and precipitation.

### Projected Shifts in Environmental Suitability of Key Malaria Vectors

To investigate the current and future environmental suitability of the six species, we fitted ensemble models using land use and climate data as covariates. Our future projections show substantial increases in environmental suitability under future climatic and land use conditions for *An. gambiae, An. coluzzii,* and *An. nili s.l.*, and minimal variation for the other three species (Fig. 2).

**Figure 2.**
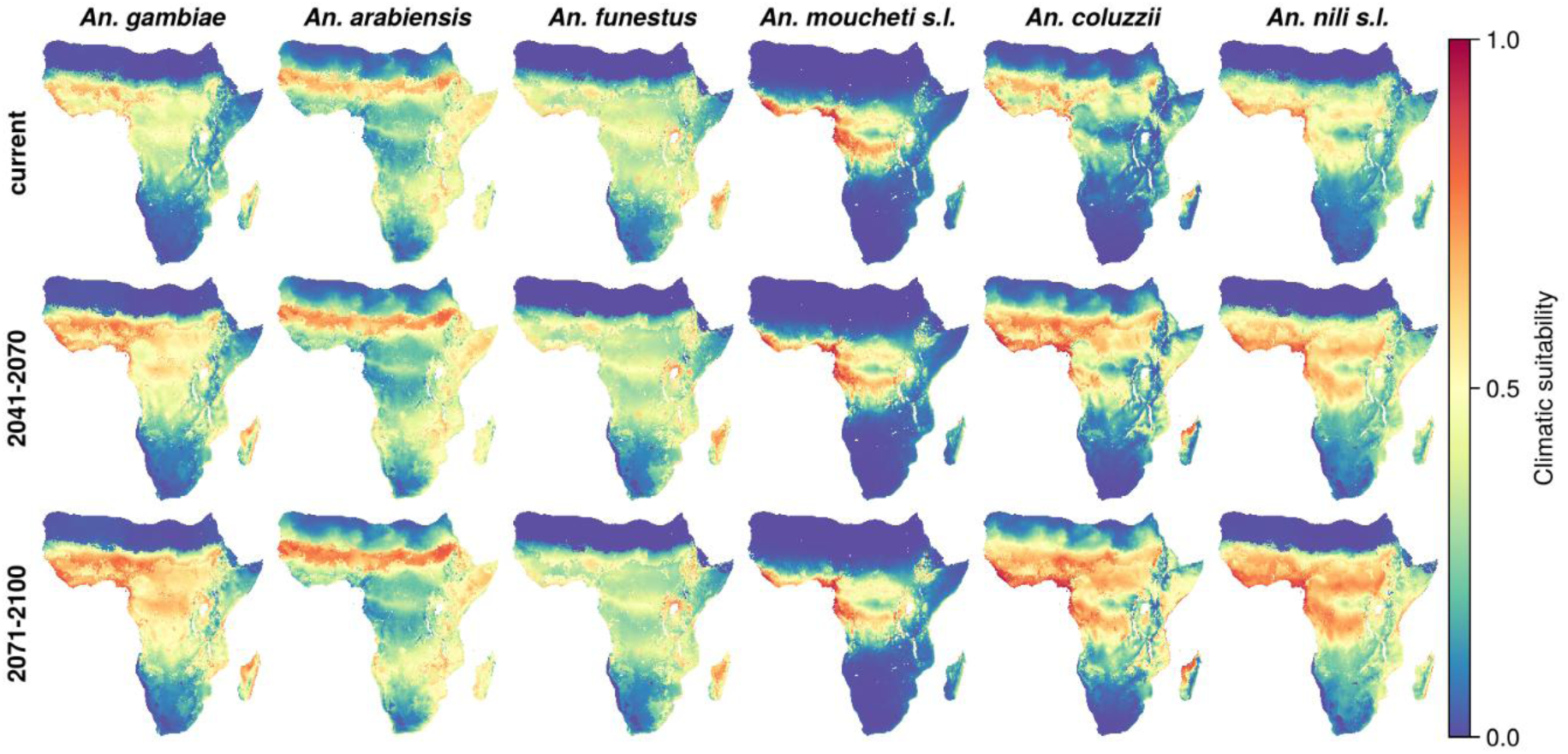
Climatic suitability of African malaria vector species under current and future conditions. Climate and land use projections shown are under the SSP3-7.0 scenario with higher emissions. Climatic suitability is projected to increase significantly for *An. gambiae*, *An. coluzzii*, and *An. nili s.l.*, whereas it remains stable for *An. arabiensis*, *An. funestus*, and *An. moucheti s.l*.

Suitable areas for *An. gambiae* are currently in West Africa, around the Great Lakes and in Mozambique whereas suitable areas for *An. coluzzii* are concentrated in West Africa and for *An. nili s.l.* in West and tropical Central Africa. For all these three species, suitability is projected to rise in already suitable areas, such as West Africa, and expand into currently less suitable regions, including in East Africa. This upward trend persists throughout the century, showing clear progression from current conditions to mid-century and end-of-century scenarios. Under the SSP1-2.6 scenario, which assumes far-reaching action to mitigate climate change, the projected increases in environmental suitability are much smaller (Extended Fig. 1).

For *An. arabiensis* and *An. funestus*, which we found are climate generalists, most of Sub-Saharan Africa is climatically suitable, except for regions in South Africa and Central Africa in the case of *An. arabiensis*. The suitable area for *An. moucheti s.l.* is restricted to tropical Central Africa (Figure 2, top row). We project that the suitable areas for these three species will remain the same under climate change (Figure 2, middle and bottom row).

The model performance of the ensembles, measured by spatially cross-validated Area Under the Curve (AUC), True Skill Statistic (TSS), and the Continuous Boyce Index (CBI), was adequate for all species (Table 1). The models for the different species had AUC values between 0.72 and 0.84, CBI values between 0.51 and 0.94.

**Table 1.** Model performance. . The average testing score is given for each metric and each species, with the score for the training data in brackets. The model performance shown was calculated from the mean prediction of the ensemble. AUC: Area Under the receiver operating Curve; TSS: True Skill Statistic; CBI: Continuous Boyce Index.

| Species | AUC | TSS | CBI |
| --- | --- | --- | --- |
| *An. gambiae* | 0.76 (0.79) | 0.44 (0.47) | 0.88 (0.99) |
| *An. arabiensis* | 0.73 (0.76) | 0.36 (0.41) | 0.94 (0.99) |
| *An. funestus* | 0.72 (0.78) | 0.38 (0.42) | 0.81 (0.97) |
| *An. moucheti s.l.* | 0.84 (0.92) | 0.63 (0.73) | 0.51 (0.96) |
| *An. coluzzii* | 0.82 (0.84) | 0.55 (0.56) | 0.82 (0.98) |
| *An. nili s.l.* | 0.72 (0.79) | 0.40 (0.46) | 0.70 (0.98) |

### Species- and Region-Specific Drivers of Malaria Vector Suitability

Understanding the environmental drivers of malaria transmission is crucial for pre-empting outbreaks. Therefore, we explored the contribution of different climatic and land-use variables controlling the distribution of mosquito species carrying malaria. Out of six predictor variables (two of which are related to temperature, three to precipitation, and one to land use), annual precipitation stood out as the first or second most important variable for five out of six investigated species (Table 2). Annual mean temperature is also a key predictor for *An. nili s.l., An. coluzzii*, and *An. gambiae*, which are also the three species projected to go through increases in suitability, while temperature annual range is important to explain the distribution of *An. moucheti s.l.*, which is restricted to tropical areas. Land use generally contributed less than the climate variables.

**Table 2.** Relative variable importance for each variable and each species. . The importance is given as the mean absolute Shapley value, which is the average size of the contribution of each variable to the suitability estimate. Bio12 (Annual Precipitation) is an important variable across all species, while bio1 (Annual Mean Temperature) is important for several species, including as *An. gambiae*. The bioclimatic variable of highest importance for the species is highlighted in bold. Variables are: bio1: Annual Mean Temperature; bio7: Temperature Annual Range: bio12: Annual Precipitation; bio14: Precipitation of Driest Month; bio15: Precipitation Seasonality; lulc: Land Use and Land Cover

| Species | bio1 | bio7 | bio12 | bio14 | bio15 | lulc |
| --- | --- | --- | --- | --- | --- | --- |
| *An. gambiae* | 0.074 | 0.015 | **0.153** | 0.020 | 0.023 | 0.055 |
| *An. arabiensis* | 0.028 | **0.111** | 0.061 | 0.056 | 0.083 | 0.064 |
| *An. funestus* | 0.013 | 0.032 | **0.092** | 0.033 | 0.047 | 0.075 |
| *An. moucheti s.l.* | 0.010 | **0.080** | 0.066 | 0.028 | 0.035 | 0.012 |
| *An. coluzzii* | **0.119** | 0.056 | 0.109 | 0.024 | 0.041 | 0.036 |
| *An. nili s.l.* | **0.076** | 0.028 | 0.066 | 0.010 | 0.040 | 0.069 |

Our results also reveal spatial patterns of variable contributions. For *An. gambiae,* precipitation variables explain the low suitability of this species in the Namibian, Somalian, and Sahara Desert (Fig. 3), with higher precipitation linked to higher suitability. Warmer areas, such as in West Africa have higher Shapley values for temperature variables, indicating that temperatures contribute to higher suitability of *An. gambiae*. “Cropland” and “Urban” land cover was consistently associated with higher suitability and Barren land with lower suitability, although land cover was a much less important predictor for *An. gambiae* suitability than climatic variables. The increasing suitability under future (SSP3-7.0, 2071-2100) conditions compared to current conditions was almost entirely explained by temperature variables (Fig. 3, bottom row).

**Figure 3.**
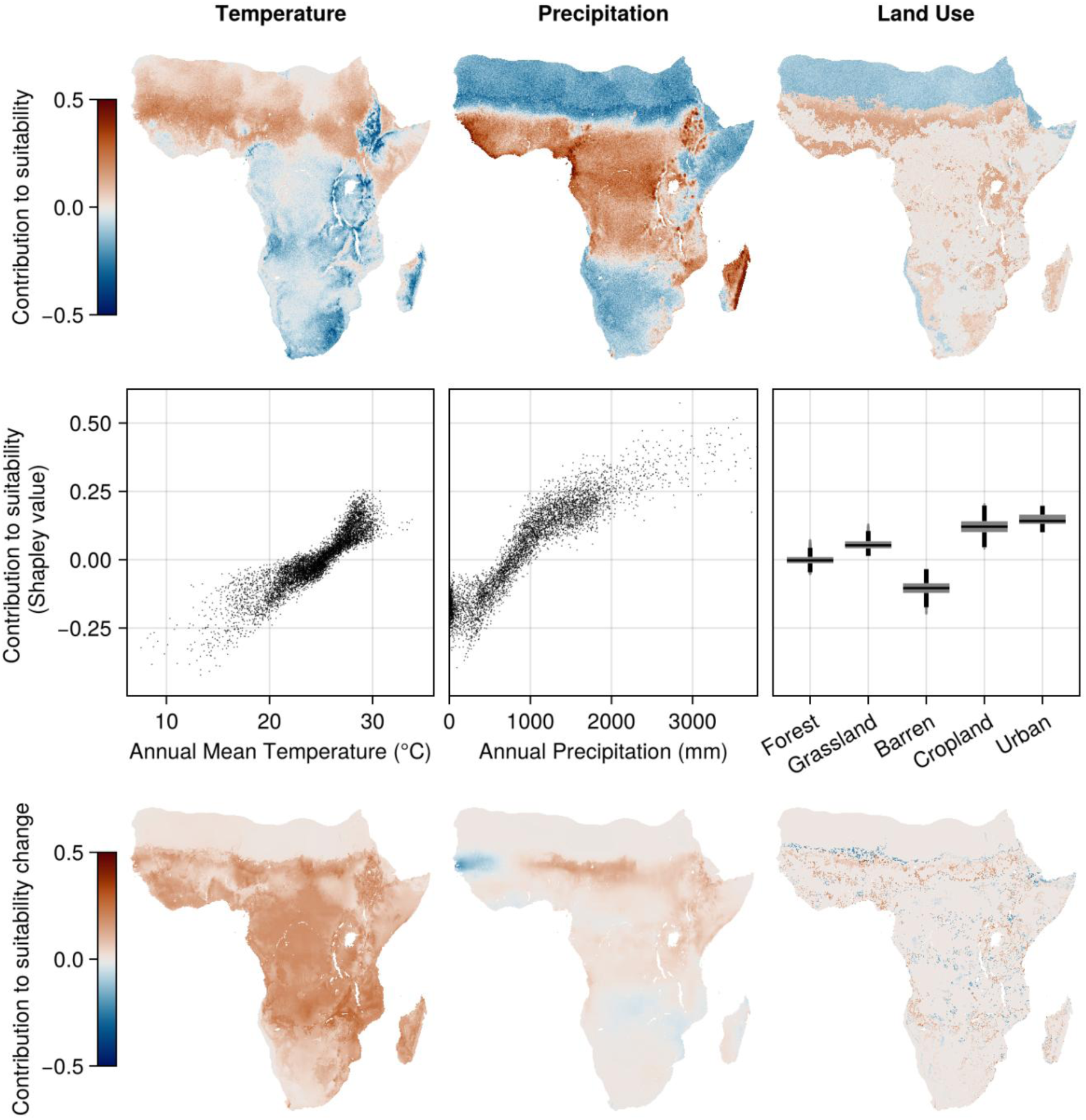
Model explanations for *Anopheles gambiae* based on Shapley values. A Shapley value measures the contribution of each variable to the final prediction for any grid cell (positive values (red) indicate the variable increased the suitability, whereas negative values indicate the variable decreased suitability, and values close to 0 indicate no contribution whatsoever. For clarity, Shapley values for temperature variables (left row) and precipitation (middle row) variables are shown combined. **Top row:** Shapley values plotted for the whole region of interest. In arid areas such as the Sahara, precipitation variables contribute negatively to suitability of *An. gambiae*. **Middle row:** Shapley values plotted against temperature, precipitation, and land use values for ten thousand randomly drawn grid cells. Higher temperature and precipitation correlate with higher Shapley values. **Bottom row**: Shapley values calculated using Baseline Shapley, demonstrating that changing temperature explains most of the increase in suitability between current and future (SSP3-7.0, 2071-2100) conditions.

Variable contributions for the other malaria vector species modelled had different geographic and environmental patterns (Extended Data Fig. 2–6). For *An. arabiensis*, precipitation variables, but not temperature variables favour suitability in arid areas (Extended Data Fig. 2). For *An. moucheti s.l.*, both temperature and precipitation variables explain the suitability in Central Africa (Extended Data Fig. 4).

### Implications of changing mosquito distributions

Modelled vector abundance has been successfully used to guide malaria interventions^36^, but the usage of vector suitability maps such as those presented here to make inference about actual malaria risk, is not necessarily straightforward. Although the presence of a competent vector is a requirement for malaria transmission to take place, many other factors, of both socio-economic, demographic and other bio-climatic nature, such as the availability of healthcare, protection against mosquito bites, and air temperatures play an important role for malaria infection risk. Furthermore, the link between environmental suitability and mosquito abundance remains uncertain^38^. To investigate the connection between vector environmental suitability and actual malaria prevalence, we analysed if vector suitability is associated with observed prevalence of childhood malaria despite these limitations. For *An. gambiae*, *An. nili s.l.*, and *An. funestus*, areas with higher suitability consistently had higher observed malaria prevalence, with the highest correlation coefficient for *An. gambiae* (r = 0.53) (Fig. 4). The association between suitability and malaria prevalence was more unclear for the other species. The differences between species might reflect the different roles these vectors play in malaria transmission, while the overall positive correlation suggests vector suitability is a useful indicator for malaria risk.

**Figure 4.**
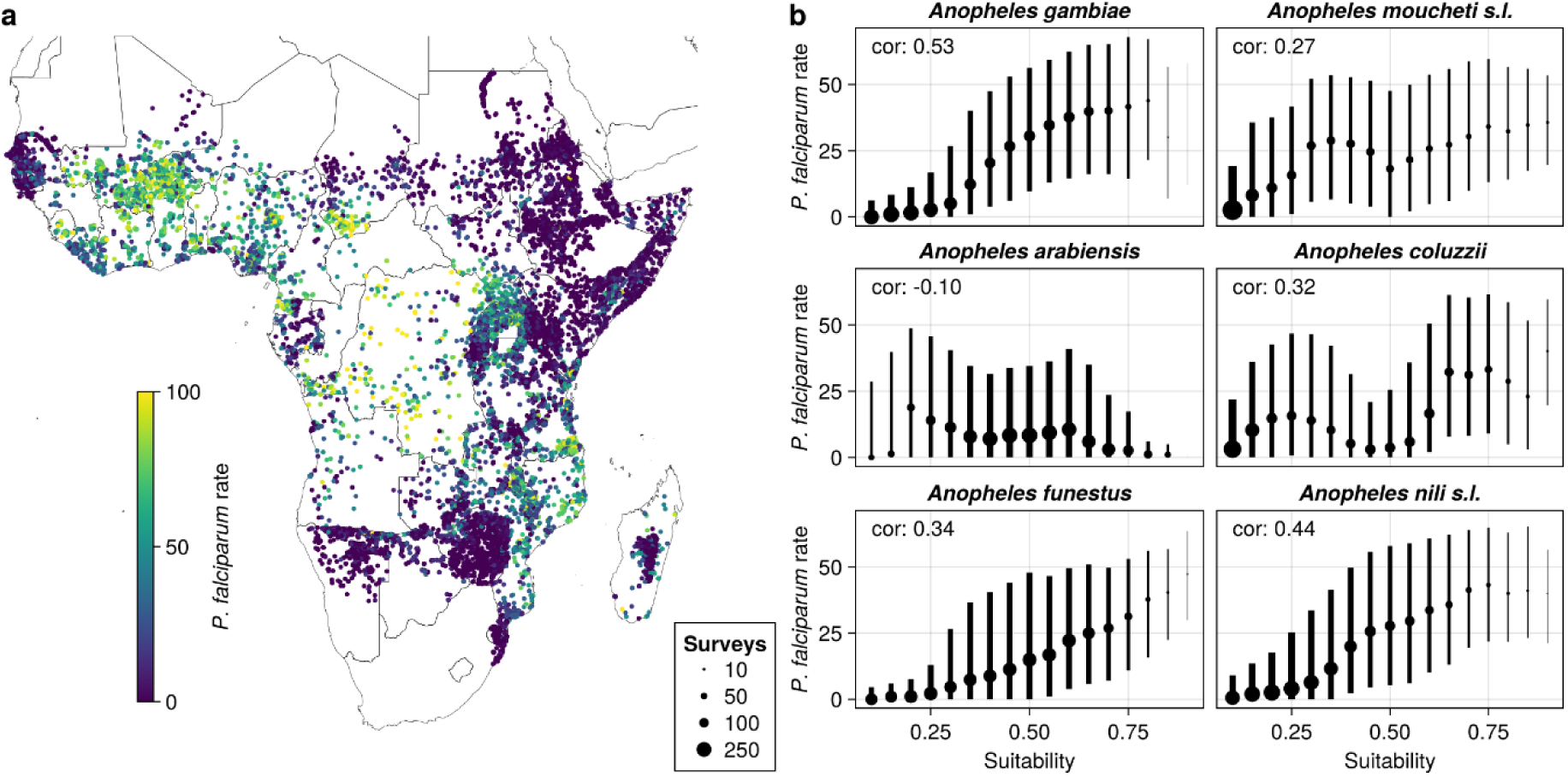
Suitability of malaria vectors is correlated with childhood malaria across Sub-Saharan Africa. **a)** Plotted survey data of childhood malaria prevalence (age-corrected *P. falciparum* prevalence in children aged 2-10) show that malaria prevalence is high in Western and Central Africa. **b)** Higher suitability of *An. gambiae*, *An. nili s.l.*, and *An. funestus* is consistently associated with higher *P. falciparum* rates, as shown by median (dots) and interquartile ranges (lines) in each bin, as well as by the correlation coefficient. For each species, surveys were divided into overlapping bins 0.2 points wide and spaced 0.05 points apart. The size of dots represents the number of surveys in each bin.

As climate and land use change continue to reshape malaria risk, identifying new emerging hotspots where high vector suitability and human population density overlap is crucial, especially as it could mean exposing vulnerable, immunologically naive populations to threats from malaria. Currently, most areas that are both densely populated and highly environmentally suitable for *An. gambiae* are found in West Africa (Fig. 5a). In the future, these areas will become even more suitable due to environmental changes. In addition, new hotspots with high human population density and high *An. gambiae* suitability will emerge in East and Central Africa, especially under the SSP3-7.0 scenario, underscoring the need for enhanced surveillance and control measures in these regions.

**Figure 5.**
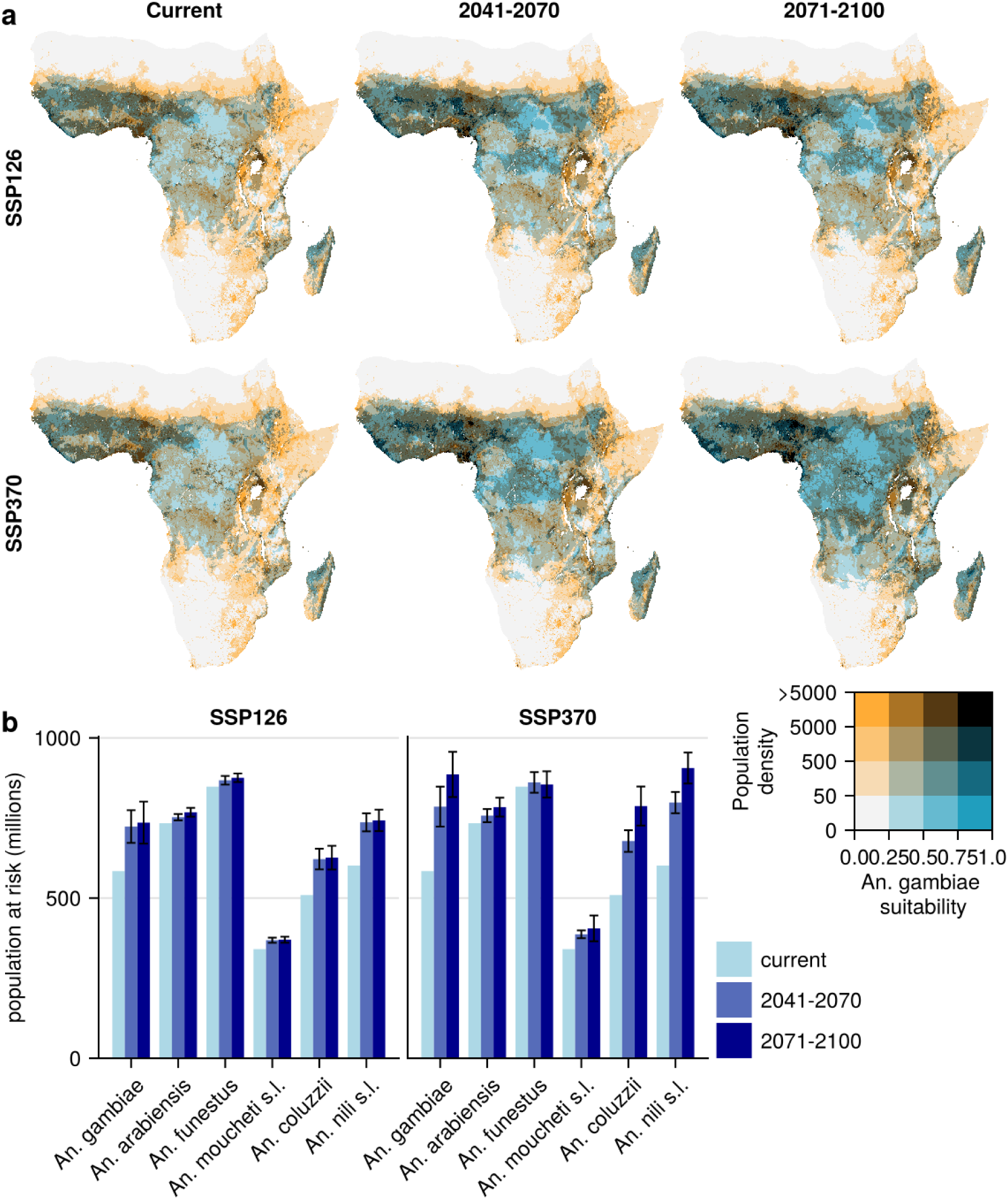
Overlaps of high climatic suitability for *Anopheles* mosquitoes and human population densities. Estimates are illustrated under current environmental conditions and future conditions, according to the low-emission SSP1-2.6 and high-emission SSP3-7.0 scenario. **a)** Human population density plotted with the suitability of *An. gambiae*, the principal malaria vector, under current (left panel) and future (middle and right panels) climate scenarios. **b)** Population that lives in areas highly suitable for each of six malaria vector species. Error bars indicate the range of the estimates across 5 climate models used, while the bars correspond to the mean estimate.

More than 800 million people currently live in areas predicted as highly suitable for *An. funestus*, the highest number for any of the investigated vectors (Fig. 5b). However, while the number of people that live in areas highly suitable for *An. funestus* as well as *An. moucheti s.l.* and *An. arabiensis* will stay stable until the end of the century, the number of people that live in areas highly suitable for *An. gambiae, An. coluzzii*, and *An. nili s.l.* will increase significantly. Under the SSP3-7.0 scenario, the areas highly suitable for *An. nili s.l.* and *An. gambiae* will respectively host 906 million (range: 863-965 million) and 885 million (range: 825-984 million) people by the end of the century, overtaking *An. funestus*. These figures are almost certainly underestimates, since we did not account for future demographic changes.

While the expansion of malaria vectors continues throughout the century under the high-emission SSP3-7.0 scenario, in the low-emission SSP1-2.6 scenario, there is almost no change between the middle and end of the century (Fig. 5). This highlights that aggressive climate mitigation can still stave off at least some of the effects of climate change.

## Discussion

We find that climate and land use change could reshape the distribution of Africa’s top malaria vectors, which could potentially redefine future malaria risk patterns in Africa. We highlight three species as particularly important for a shifting landscape of malaria risk across Africa: *An. gambiae*, *An. coluzzii,* and An. *nili s.l.* Due to climate and land use change, the number of people living in areas highly suitable for these species could increase by some 200 million people, with new hotspots emerging in East and Central Africa and continued high suitability in West Africa (Fig. 5). These findings highlight the urgent need to adapt vector control strategies to shifting vector distributions driven by climate change, ensuring interventions remain effective in regions where malaria risk is likely to intensify.

The predicted changes in environmental suitability of malaria vectors could present a challenge to malaria control. If increasing environmental suitability would trigger higher mosquito abundance^38^, this directly increases the force of infection of malaria^10,39,40^. Moreover, if malaria vectors emerge in new areas, this can challenge existing control interventions that do not target the newly emerging species^41,42^. The (re-)emergence of vectors could especially present challenges in areas where malaria transmission is currently low, including in East Africa. For example, a large resurgence of malaria in Rwanda in 2017 has been linked to an increased number of *An. gambiae* mosquitoes and above-average temperatures^43^. While continuous control programmes across Africa have brought malaria prevalence down significantly already^44^, and ambitious goals have been set for future malaria eradication^45^, the increasing environmental suitability of key malaria vectors could challenge their continued effectiveness.

Our projections of expanding environmental suitability for key malaria vectors (Fig. 2) align with observed range shifts of *Anopheles* species. A recent analysis of the historical occurrence records of 21 *Anopheles* species showed that these have been recorded at increasingly polewards and upslope locations in the course of the 20^th^ century^30^. Examples of newly established vectors contributing to malaria transmission include the introduction of *An. gambiae* after rainforest clearance in Cameroon^26^, the emergence of *An. arabiensis* in Guinea-Bissau^46^, and the involvement of the invasive malaria vector *Anopheles stephensi* in urban malaria^47^. However, since factors such as vector control efforts and the increasing human footprint^48^ may also contribute to shifts in vector distribution and malaria transmission, it is difficult to attribute any particular change to climate change^49,50^. To understand how these processes play out locally and globally, there is a continued need for longitudinal field studies that can detect the changing distribution of malaria vectors.

Our finding that *Anopheles* vector mosquitoes are favoured by climate change is consistent with a recent SDM study on *Anopheles* mosquitoes^51^ and our current-day projections generally align well with the distribution maps generated by the Malaria Atlas Project^23,27^. However, this finding stands in contrast with earlier projections for *An. gambiae* that were published over 10 years ago^52–55^. This might reflect advancements in data availability and SDM methods since these studies were published. Furthermore, our models project that the eastern Sahel region, including South Sudan and western Ethiopia, are moderately suitable for *An. gambiae*, despite the absence of records from this area in our dataset. A survey that is not in our dataset indicates that *An. gambiae* is absent in Sudan, but present in South Sudan^56^. This illustrates remaining issues with sampling bias and sparsely sampled regions, where more research is needed to ascertain the actual distributions of malaria vectors.

Mechanistic models have been instrumental in advancing our understanding of malaria transmission by providing valuable insights into the thermal limits of parasite development and vector survival^12,19,57–61^. Mordecai et al.’s influential paper on malaria transmission by *An. gambiae* concludes that malaria transmission peaks around 25°C^3^, in part due to decreased mosquito survival and reproduction above this temperature^62–64^. Numerous follow-up studies predicted a decrease in malaria transmission across most of sub-Saharan Africa, and especially in West Africa^5,13,18,19,59,65^. However, this stands in contrast with our findings that the environmental suitability of malaria vectors including *An. gambiae* will increase across sub-Saharan Africa with climate change.

Possible explanations for this discrepancy reflect the inherent limitations of the respective modelling approaches used. In mechanistic modelling studies, it is hard to account for behavioural adaptations that allow species to survive in adverse conditions^66^. *Anopheles* mosquitoes have evolved a number of behavioural adaptations to persist in adverse conditions, including aestivation during the dry season, seasonal recolonisation, and resting and feeding indoors^67,68^. Genetic plasticity may also play a role in adaptation to local conditions^69,70^. On the other hand, regardless of adaptations, all species must have an upper thermal limit, even if it cannot be observed from current distribution patterns^71^. The uncertainty connected with extrapolation to novel climates is a crucial limitation of SDMs, particularly when no upper limit can be detected from occurrence data.

We did not find a clear upper thermal limit for several species including *An. gambiae*, which is consistent with another recent study that used SDMs to estimate the thermal limits of 7 mosquito species failed to find upper limits for five of those, including *An. gambiae*^72^. This increases the uncertainty of predictions of mosquito distributions towards future climates.

We also show that precipitation, not temperature, variables are the most important predictors for most of the investigated species (Table 2). For example, we find that *An. gambiae* is associated with higher precipitation and that precipitation limits suitability of *An. gambiae* in the Sahel region (Fig. 3). In contrast, arid conditions are suitable for its sibling species *An. arabiensis,* which is consistent with earlier knowledge about the ecology of these species^73,74^. While the role of water availability in malaria transmission has been recognized before^20,75–77^, these studies do not link it to the distribution of individual vector species. Further investigations into the species-specific responses to variations in precipitation could be key to inform vector control strategies and improve predictive models.

Since we limited this study to malaria vectors native to Sub-Saharan Africa, we have not investigated *Anopheles stephensi*, an invasive species that is spreading quickly in Sub-Saharan Africa and is a highly anthropophilic and urban species^78–80^. It was first observed in Djibouti in 2012 and has since spread to Ethiopia, Sudan, Somalia, and Kenya^78,81^, where it is already involved in malaria transmission^47^. Multiple studies have already investigated the potential spread of *An. stephensi*, showing that this species might spread to numerous urban areas in Africa and representing yet another challenge on the path to malaria eradication^82,83^.

Recent reviews have highlighted shortcomings in SDM studies on mosquitoes^29,84^. Our study addressed some of these by using ensemble modelling, using occurrence data from the full range of species, and carefully selecting predictor variables^85^. However, several limitations remain, such as the reliance on data collected from a variety of literature sources and varying quality, the paucity of occurrence records in Central Africa, and the uncertainty associated with projecting models into no-analogue climates^86,87^. Our study also does not capture all factors influencing mosquito distribution, such as direct human interaction and habitat modifications^88,89^. While we obtained an acceptable to good model performance for most species, some anophelines have a very wide distribution across varied habitats and might therefore be inherently challenging to model^90^.

Malaria remains the deadliest vector-borne disease on earth, and any impact of climate change will have huge ramifications, in particular in sub-Saharan Africa. We project that the suitability for three key malaria vector species will increase with climate change across sub-Saharan Africa. Given the enormous health importance of these mosquitoes, additional research that addresses the ongoing redistribution of malaria vectors is needed, including longitudinal field studies, experimental work on less-studied vector species^91^, and modelling that integrates mechanistic and correlative methods to reconcile laboratory observations of the thermal performance of *Anopheles* species with their observed distribution and observed malaria prevalence. As the world strives for malaria elimination while dealing with accelerating climate change^34,35,45^, we stress the need to keep on-going control and eradication efforts efficient in the face of the anticipated distribution changes of malaria vectors to keep this target within reach.

## Methods

### Mosquito occurrence data

Based on earlier reviews, Sinka et al. propose seven dominant vector species in Sub-Saharan Africa: *Anopheles gambiae*, *Anopheles arabiensis*, *Anopheles funestus s.s.*, *Anopheles arabiensis, Anopheles merus, Anopheles melas, Anopheles nili s.l.*, and *Anopheles moucheti s.l.*^23^. Of these, *An. merus* and *An. melas* are saltwater species with restricted to coastal regions, and we therefore excluded them from this study. Furthermore, in 2013 the M-form of *An. gambiae* was recognized as a separate species and has since been known as *Anopheles coluzzii*^92^. Both *An. gambiae* and *An. coluzzii* are highly efficient malaria vectors, with distinct behaviours and environmental preferences. We therefore model both separately. In total, we thus model six *Anopheles* species that all can act as primary malaria vectors: *An. gambiae, An. coluzzii, An. arabiensis, An. moucheti s.l., An. nili s.l,* and *An. funestus*.

Mosquito occurrence data was extracted from a database published by Kyalo et al. in 2017, which is one of the most extensive databases on *Anopheles* species, including more than a century of records of 21 species^22,93^, and the database collated by the Malaria Atlas Project, which was accessed both through a published dataset^94,95^ and the malariaAtlas R package^96^.

The databases were collated, different synonymous species designations were converted as appropriate (e.g. *An. gambiae s.s. M form* to *An. coluzzii*) and duplicate records removed. We limited the data to the period between 1980 and 2010 in order to match the period of the 30-year average of the climate data. We used spatial thinning in order to reduce spatial bias, employing a thinning distance of 10 kilometres^97^. This resulted in between 166 (for *An. moucheti s.l.*) and 2773 (for *An. arabiensis*) records for each species (Fig. 1)

### Background data

Occurrence data often has spatial bias, resulting from unevenly distributed sampling efforts. It is challenging to determine whether spatial patterns are a result of sampling bias or differences in habitat suitability. Different strategies for generating background points range from sampling randomly in geographic space, sampling in environmental space, or using presence records of similar species as background points^98,99^. Here, we choose a middle-of-the-road approach, sampling 50% of background points randomly within the region of interest, and 50% from sampling locations where other anopheline species were found. The total number of background points generated was twice the number of presences.

Since the species we modelled are all distributed throughout Sub-Saharan Africa, but none of them are found north of the Sahara, we defined our region of interest based on the Sahara Desert and drew background points only from this region. The northern boundary of our region of interest was defined by taking the boundary where precipitation is at 200 mm per year and buffering this by 7 degrees (approx. 750 kilometres).

### Climate and land use data

We used climate data from CHELSA, which provides bias-corrected climate variables with global coverage at a 1 km spatial resolution^100^. For future projections, we used projections for the mid-century (2041-2070) and end of the century (2071-2100), under a low (SSP1-2.6) and a high (SSP3-7.0) emissions scenario for each of the five global circulation models (GCMs) available through the CHELSA website. We selected two temperature and three precipitation variables that we deemed biologically relevant for our species in question and were not strongly correlated in the region of interest to avoid multicollinearity issues (Pearson correlation coefficient <0.7 for all variables). These were bio1 (Annual Mean Temperature), bio7 (Temperature Annual Range), bio12 (Annual Mean Precipitation), bio14 (Precipitation of Driest Month), and bio15 (Precipitation Seasonality). Data were aggregated to a resolution of 2.5 arcminutes (ca. 5 kilometres).

In addition to climate data, we used a global land use product with 7 land cover features^101^. This dataset is also available under the SSP1-2.6 and SSP3-7.0 scenario. For the mid-century, the projection for 2055 was used and for the end of the century the projection for 2085. The data was resampled to match the resolution and projection of the climate data, and any cells classified as “Water” were masked.

### Ensemble modelling

Ensemble modelling involves fitting multiple machine learning models to the same dataset and combining their outputs. This approach is widely used in SDMs to mitigate algorithmic uncertainty^102^. Here, we used an ensemble with three models: a generalized additive model (GAM), Maxnet (a close analogue to Maxent)^103^, and an artificial neural network (ANN). These models were selected for their ability to capture complex interactions between variables without being too prone to overfitting, and we tweaked model settings to further reduce the risk of overfitting. We limited the flexibility of the GAMs by setting the *k*-parameter of the smooth term to 5 and the gamma parameter to 3. For Maxnet, the feature classes were limited to linear, quadratic, and product (excluding hinge and threshold features). For the ANNs, we used only a single hidden layer with 16 neurons and fit for 50 epochs.

Modelling was done in Julia v1.11, a high-performance scientific computing programming language, using the MLJ machine learning framework^104^. For GAMs, we used the *mgcv* library in R^105^, interfacing between the two programming languages with the Julia package RCall. We used Julia’s Flux.jl package to fit ANNs^106,107^ and Maxnet.jl to fit Maxnet models.

Spatial autocorrelation can inflate performance metrices for models using spatial data, as has often been observed in SDMs^108,109^. Here, we use 5-fold spatial cross-validation to more accurately measure the performance of the ensemble model. The study region was divided into squares of 3 by 3 degrees (approx. 300 by 300 kilometres), which were then randomly distributed into five folds. The ensemble was fit five times, leaving out one fold for evaluation each time. To assess the performance of the model, we used the Area under the Receiver operating characteristic Curve (AUC), the True Skill Statistic (TSS), and the Continuous Boyce Index (CBI)^110,111^. These metrics capture different aspects of model performance; whereas AUC and TSS measure the ability of the model to discriminate between presence and absence points, the CBI is a presence-only metric that assesses if areas with higher suitability scores consistently contain more occurrence records. We report the average of each performance metric over the five folds.

After model validation, ensembles were fit using all data available for each species. These ensembles were then used to generate projections to current and future climates. The final prediction is the simple mean of the prediction of each model in the ensemble. For predictions under future climate conditions, predictions were generated for each of five GCMs and then averaged.

### Model interpretations

We used Shapley values to understand how each variable contributes to the ensemble prediction and their relative importance. The Shapley value is a game theory concept that has been adopted to assess variable contributions in machine learning models^112,113^, including SDMs^33,114,115^. The Shapley value quantifies the contribution of each variable to a prediction and can be obtained by averaging the change in outcome when adding all variables in all possible orders. Shapley values have several desirable features, such as effectively handling multicollinearity and synergistic effects^116^. The sum of Shapley values of all variables is equal to the difference between the expected outcome and model outcome.

To understand how each variable contributes to the suitability under current conditions, we use a Monte Carlo algorithm^113^. This algorithm works by substituting variables in each grid cell with variables from a random other grid cell in random order and observing the resulting change in suitability. We performed 32 Monte Carlo samples for each grid cell and each variable. The variable importance of each variable was calculated as the mean absolute Shapley value of the resulting grid.

Then, to understand which variables drive the change in suitability between current and future environmental conditions, we used a related algorithm called Baseline Shapley, where input variables are instead compared with some self-defined baseline^117^. Here, we define current environmental conditions as the baseline and future conditions as the input and calculate the mean change in suitability for each variable as future conditions are substituted for the current conditions in every possible order.

### Correlation with malaria prevalence

To investigate how malaria vector suitability is associated with malaria prevalence, we use a recently published compendium of over 50 thousand malaria prevalence surveys collated by Snow et al.^118^. The dataset includes an age-corrected childhood malaria rate (estimated of *P. falciparum* positivity rate for children 2-10 years old) for each survey. For our analysis, we only consider surveys recorded between 1980 and 2010, matching the considered climate period. In addition, when multiple surveys were recorded within the same grid cell, we used the mean positivity rates of the surveys conducted. This resulted in childhood malaria estimates for 13,070 unique grid cells.

To visualize how *P. falciparum* prevalence changes with the suitability of each vector species, we assign each grid cell with survey data to one or more overlapping bins 0.2 suitability units wide and spaced 0.05 units apart. We then plot the median value and interquartile range of each bin. In addition, we calculate the Pearson’s correlation coefficient between *P. falciparum* rate and the modelled suitability in each grid cell for of each vector species. Because spatial autocorrelation increases the uncertainty of correlation estimates^119,120^, we here only report the correlation coefficient and refrain from making any significance claims.

### Populations living in areas highly suitable for malaria vectors

To assess the number of people potentially impacted by the changes in the distribution of *Anopheles* species, we combined the mosquito suitability maps with global gridded 1km population data available from WorldPop^121,122^. Firstly, we identified areas with high suitability for each vector species using a cut-off corresponding to the 10^th^ percentile training presence^109^. This cut-off was chosen to make it easier to compare estimates between species, and to exclude areas with few occurrence records. We estimate the number of people living in areas with a highly suitable climate for a particular vector species by taking the sum of the population counts of all grid cells where the suitability was above this cut-off. For future predictions, we calculate this for each GCM and then calculate the mean and standard deviation across estimates. We assume no change in population counts to isolate the effects of climate change.

## Data and code availability

No original data was used in this article. All data used can be freely accessed at other sources. All code used in this article and instructions on how to download the data and reproduce the analysis is available at https://github.com/tiemvanderdeure/AfricanMalariaVectors.

## Acknowledgements

This project has received funding from the European Union’s Horizon 2020 research and innovation program under grant agreement No. 101000365 (PREPARE4VBD). Anna-Sofie Stensgaard is grateful to the Knud Højgaard Foundation for supporting the Platform for Disease Ecology, Health and Climate (grant number: 16-11-1898 and 20-11-0483).

## Extended Data Figures

**Extended Data Figure 1.**
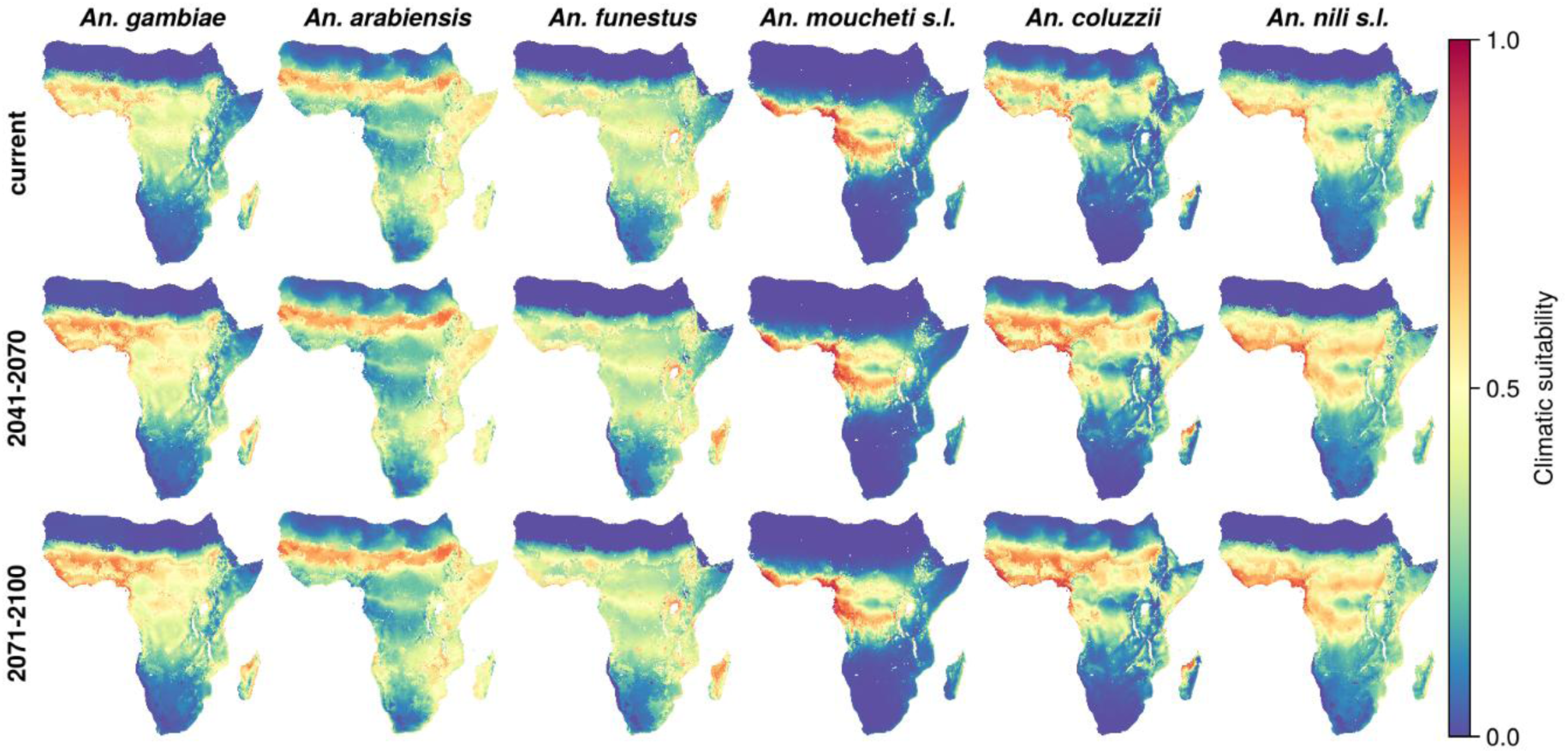
Climatic suitability of African malaria vector species under current and future conditions under the SSP1-26 scenario with low emissions.

**Extended Data Figure 2.**
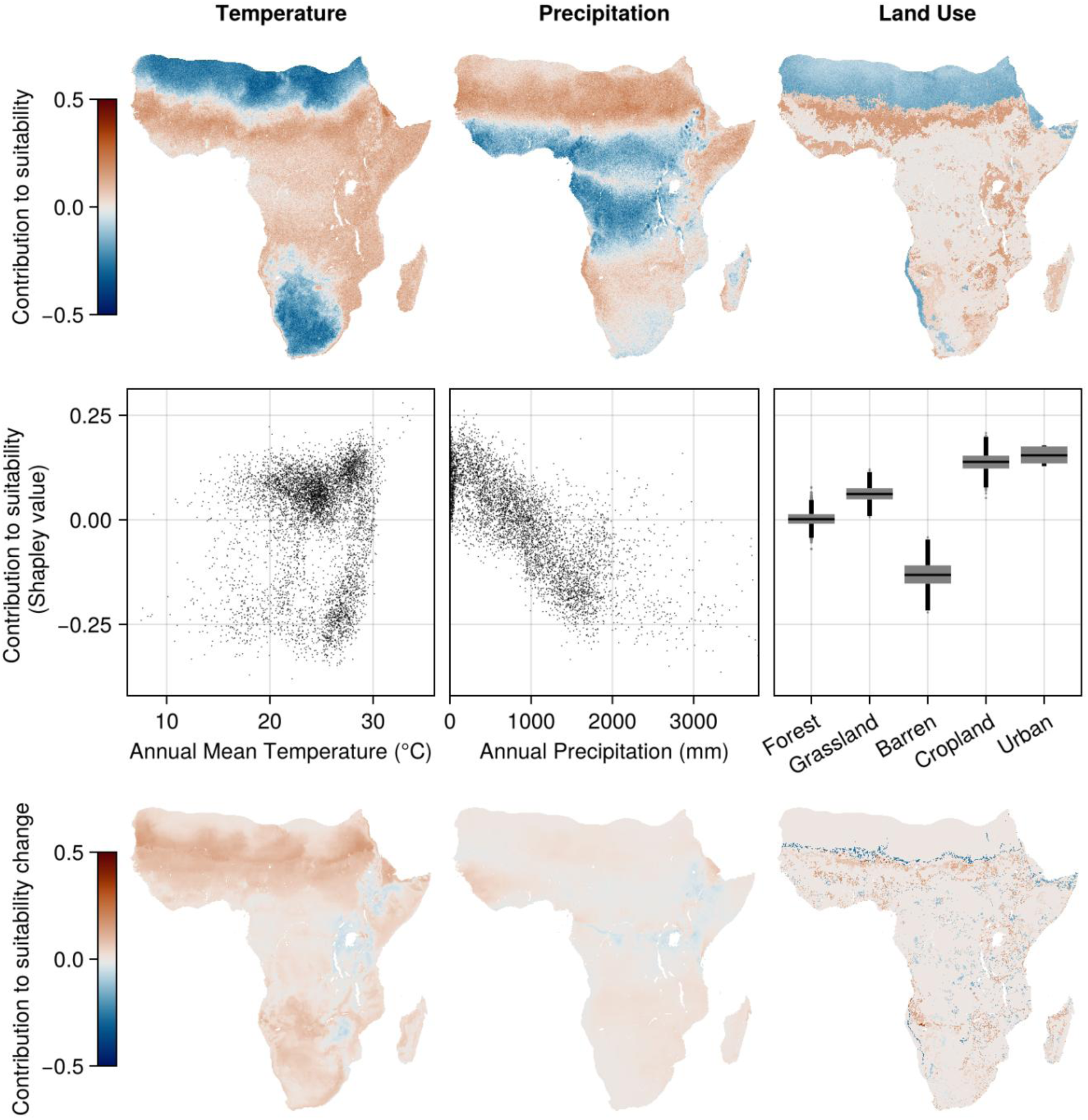
Model interpretations for *Anopheles arabiensis*.

**Extended Data Figure 3.**
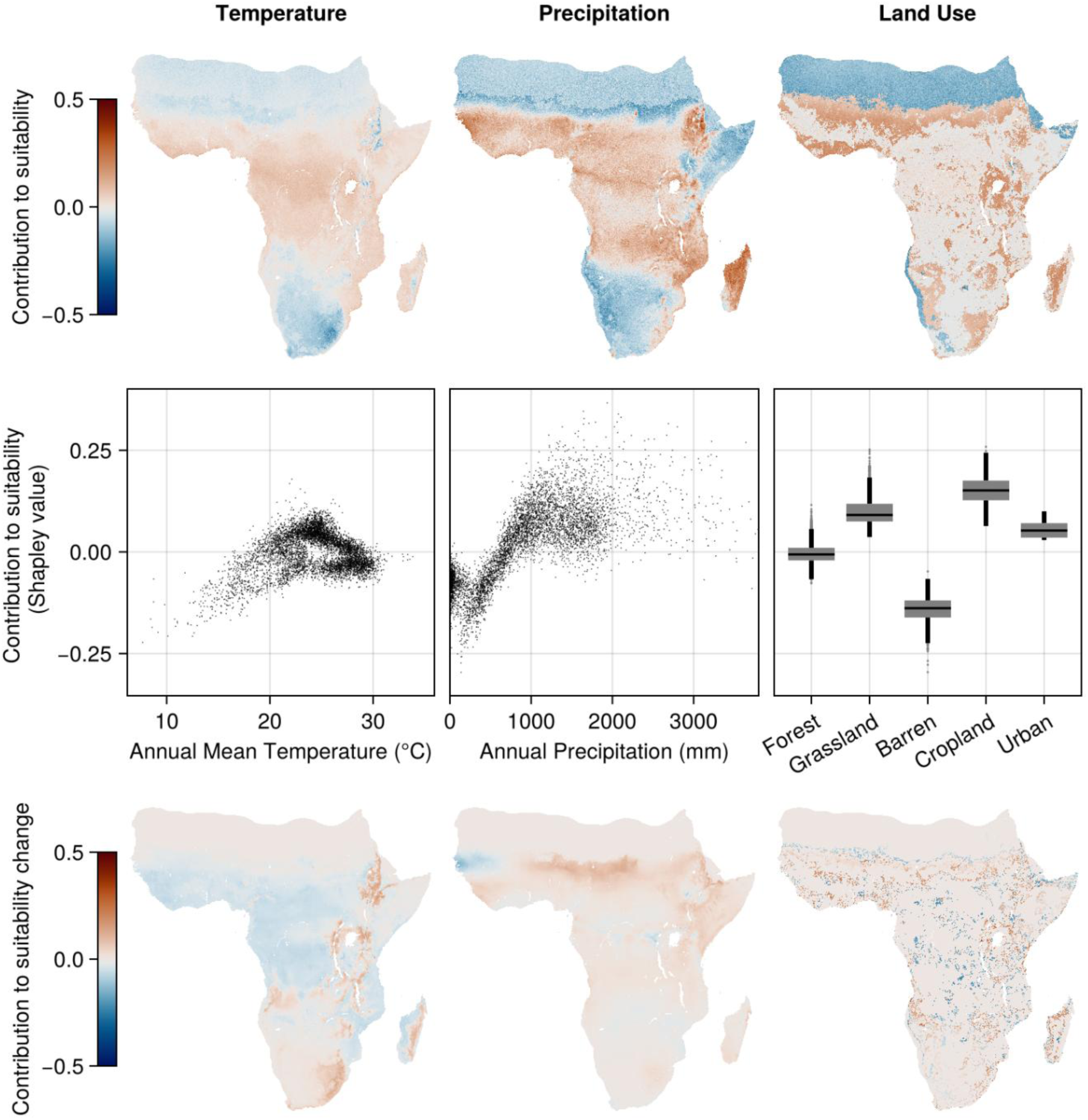
Model interpretations for *Anopheles funestus*.

**Extended Data Figure 4.**
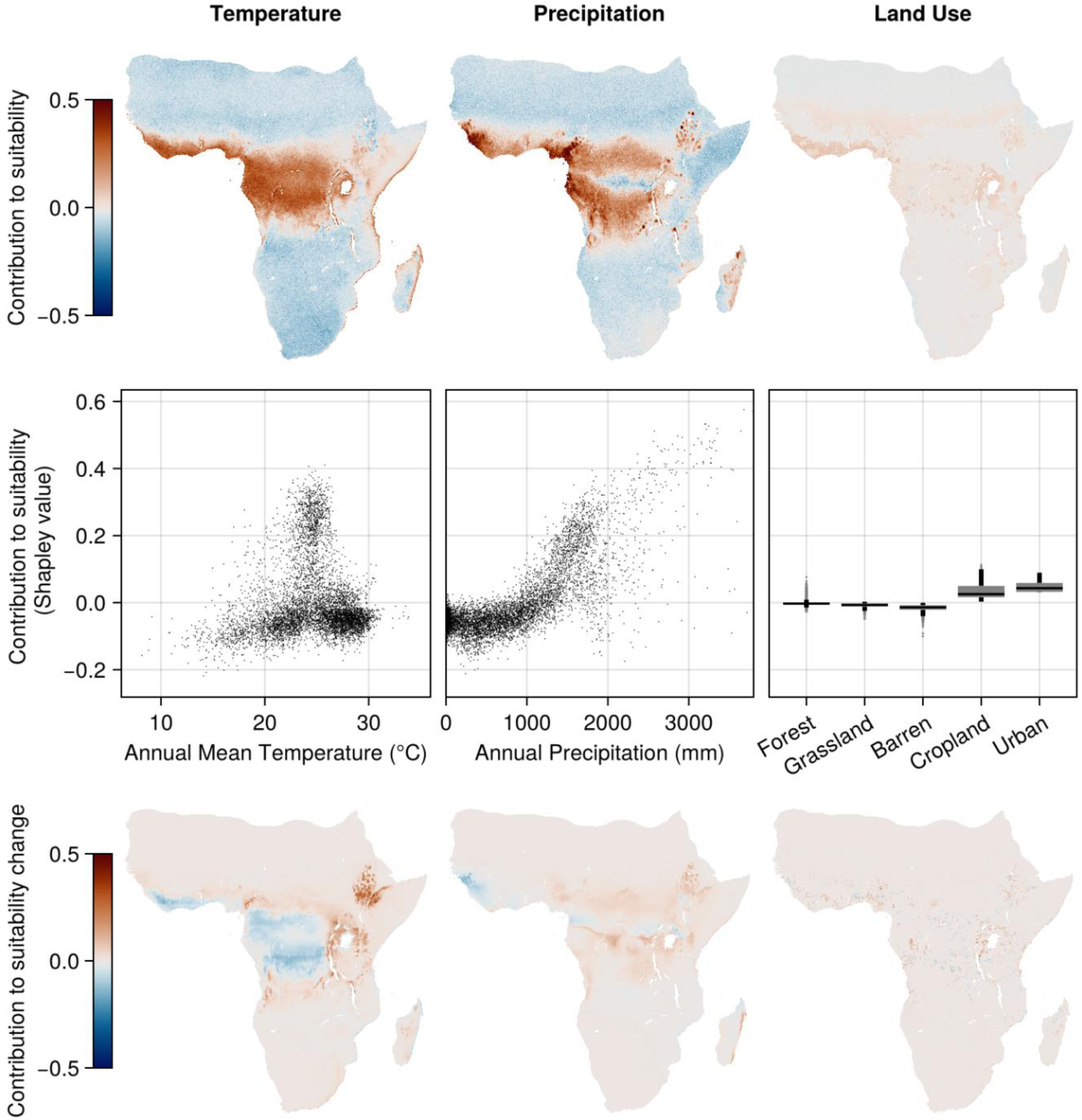
Model interpretations for *Anopheles moucheti s.l*.

**Extended Data Figure 5.**
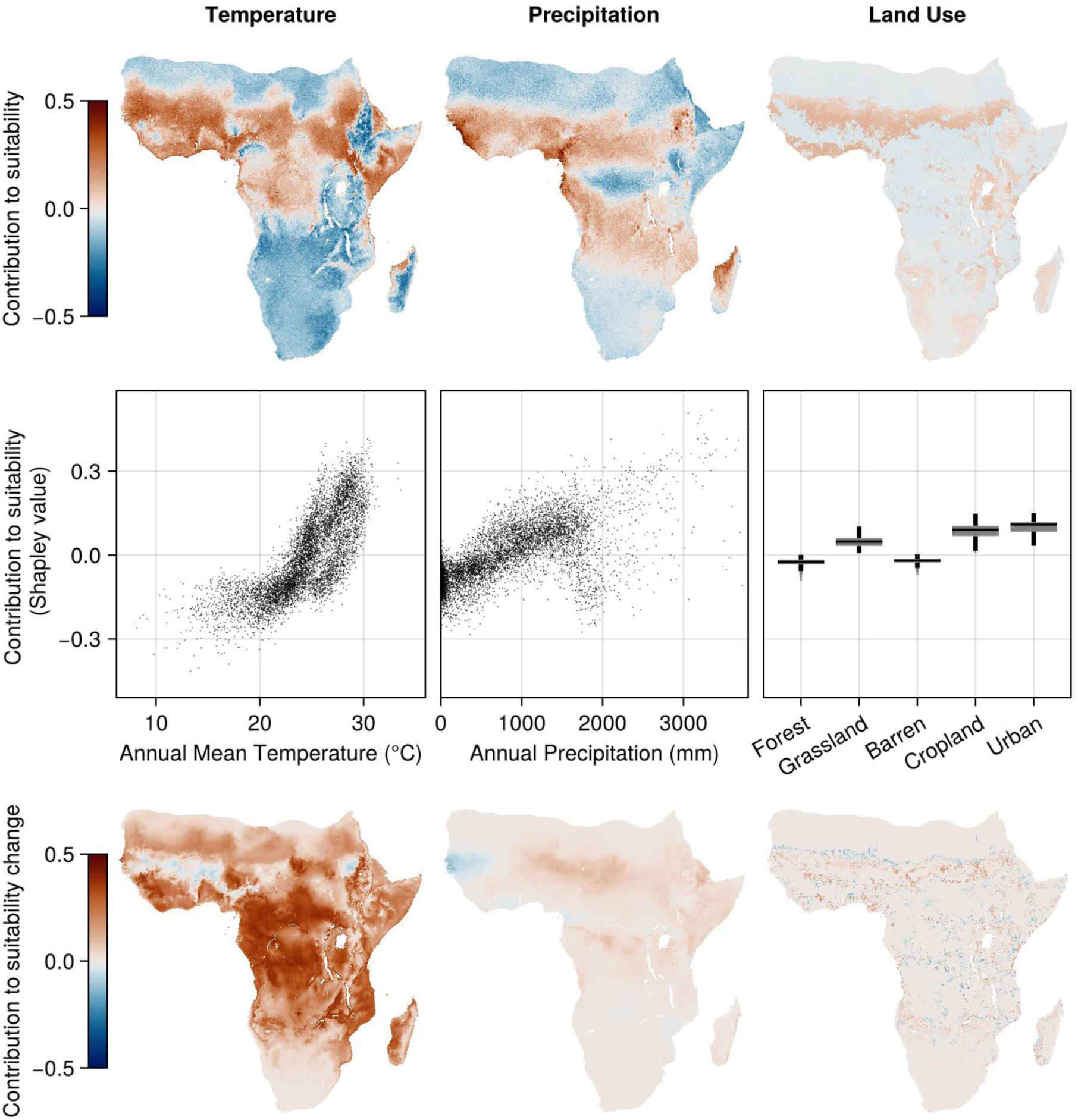
Model interpretations for *Anopheles coluzzii*.

**Extended Data Figure 6.**
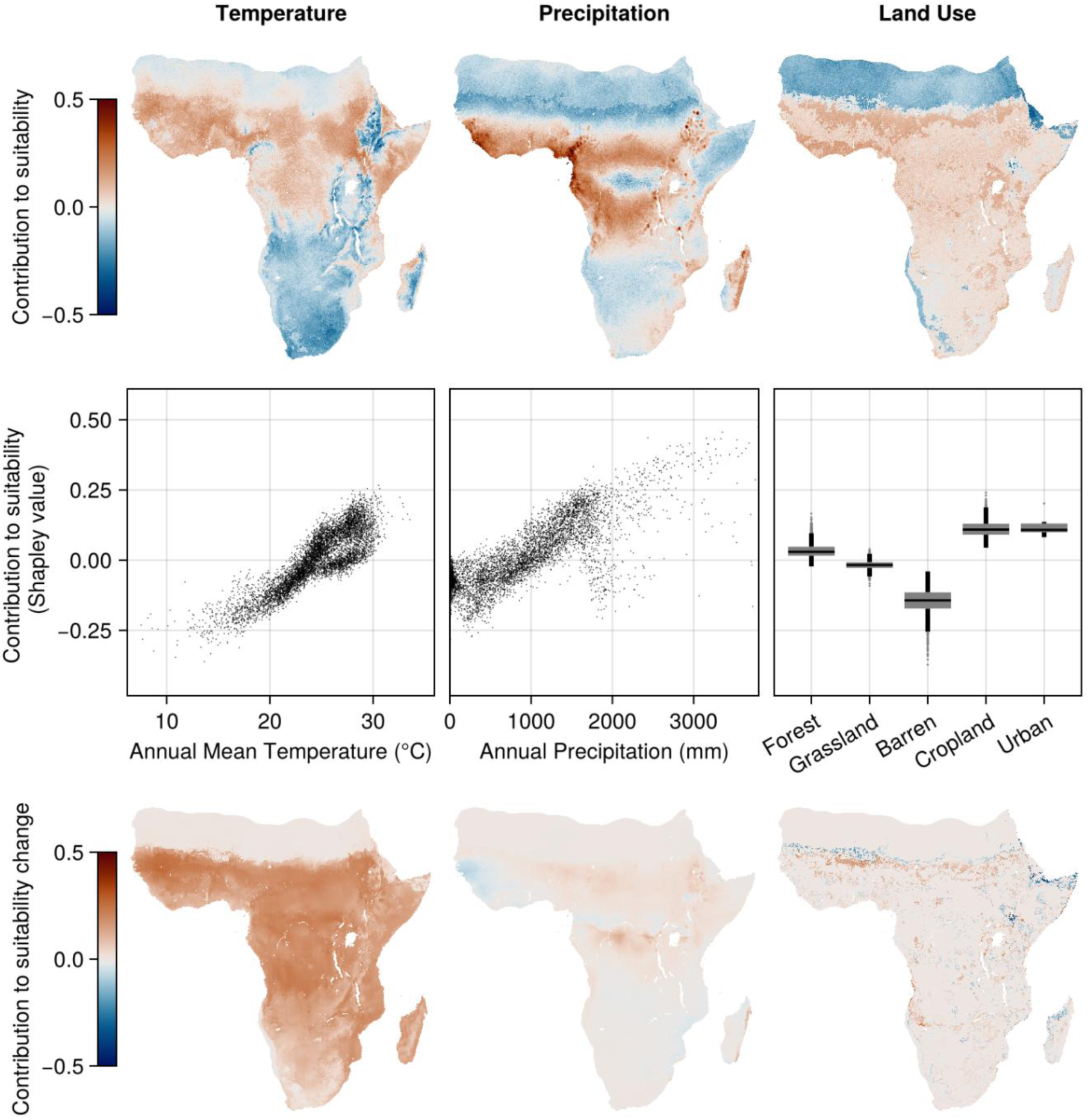
Model interpretations for *Anopheles nili s.l*.

